# Immune cell single-cell RNA sequencing analyses link an age-associated T cell subset to symptomatic benign prostatic hyperplasia

**DOI:** 10.1101/2024.12.30.629739

**Authors:** Meaghan M. Broman, Nadia A. Lanman, Renee E. Vickman, Gregory M. Cresswell, Harish Kothandaraman, Andree Kolliegbo, Juan Sebastian Paez Paez, Alexander P. Glaser, Brian T. Helfand, Gervaise Henry, Douglas W. Strand, Omar E. Franco, Susan E. Crawford, Simon W. Hayward, Timothy L. Ratliff

## Abstract

Benign prostatic hyperplasia (BPH) is among the most common age-associated diseases in men; however, the contribution of age-related changes in immune cells to BPH is not clear. The current study determined that an age-associated CD8^+^ T cell subset (Taa) with high Granzyme K (*GZMK^hi^*) and low Granzyme B (*GZMB^low^*) gene expression infiltrate aged human prostates and positively correlate with International Prostate Symptom Score (IPSS). A velocity analysis indicated that CD8^+^ T cell differentiation is altered in large BPH prostates compared to small age-matched prostates, favoring Taa accumulation. In vitro granzyme K treatment of human BPH patient-derived large prostate fibroblasts increased secretion of pro-inflammatory senescence-associated secretory phenotype (SASP)-associated cytokines. These data suggest that granzyme K-mediated stimulation of prostate stromal fibroblast SASP cytokine and chemokine production promotes prostate immune cell recruitment and activation. Overall, these results connect symptomatic BPH with immune aging.

## Introduction

Aging is associated with changes in both innate and adaptive immunity collectively referred to as immunosenescence, involving alterations in leukocyte function and increase, an individual’s susceptibility to a variety of chronic diseases, infections, and autoimmunity^1–3^. While immunosenescence encompases both adaptive and innate immunity, alterations in the adaptive immune system are a key component in the dysregulated immune response in aged individuals^4,5^. Immunosenescence includes a chronic low-grade inflammation, referred to as inflammaging, that occurs in the absence of pathogens and has been implicated in a variety of age-associated chronic conditions including cardiovascular disease, type 2 diabetes mellitus, obesity, and cancer^1,6–9^. In addition to changes in immune cells, various other cell types and cellular processes may contribute to the chronic inflammation of inflammaging, including the presence of senescent cells in the microenvironment^1,2,10,11^. Characteristics of senescent cells include cell cycle arrest, resistance to apoptosis, and a senescence-associated secretory phenotype (SASP), characterized by the production of various cytokines, chemokines and growth factors that promote an inflammatory microenvironment^2,12^. Reduced removal and accumulation of these senescent cells with age is one consequence of immunosenecence^13^.

Among the most common age-related conditions in men is benign prostatic hyperplasia (BPH), affecting approximately half of men over age 50 and nearly 80% by age 80^14–16^. BPH is characterized by progressive nodular expansion of both the epithelial and fibromuscular stromal tissues of the prostatic transitional zone (TZ) adjacent to the urethra, resulting in lower urinary tract symptoms (LUTS)^14–18^. Immune cell infiltrates that progressively accumulate over time are commonly observed in association with BPH nodules and, similar to other age-associated inflammatory conditions, are typically not associated with a pathogen^19^.

Previous studies have associated high prostate inflammatory cell infiltration with increased International Prostate Symptom Score (IPSS) and LUTS^15,20^. Also, prostatitis has been associated with a higher risk of developing BPH-associated LUTS as well as with an increased likelihood of eventually requiring medical or surgical treatment to manage these symptoms^21^. These findings led to the hypothesis that age-associated alterations in the prostate immune microenvironment may underlie the development and progression of clinical symptoms of BPH^22–24^. Immune cell-derived proteins, namely cytokines and chemokines, have been shown to promote stromal and epithelial cell activation and proliferation as well as exacerbate immune cell recruitment^22,25,26^.

Additionally, granzymes, serine proteases produced and released by cytotoxic T cells and natural killer (NK) cells upon recognition of cellular targets to exert their cytotoxic effects, have been shown to promote pro-inflammatory cytokine production by fibroblasts and other cell types^27–31^. However, the potential contributions of specific immune cell populations and their cellular products to BPH symptoms and progression have not been defined^19,20,24,32^.

Given the T cell dominance of inflammation in the aged prostate^15,33,34^, we hypothesized that age-associated T cell subpopulations contribute to BPH symptoms. To address this hypothesis, we utilized single-cell RNA-sequencing (scRNA-Seq) to define and compare the immune cells between aged and young normal prostates and between large and small age-matched prostates to identify potential age-related immune mechanisms of BPH. The results of this study suggest that an age-associated CD8^+^*GZMK*^hi^*GZMB*^low^ T cell (Taa) subset contributes to BPH symptoms.

## Results

### A CD8^+^GZMK^hi^GZMB^low^ Taa subset is present in aged prostates

Published scRNA-Seq data^33,34^ generated from immune cells isolated from TZ tissues collected from 10 “small” (≤40 grams) prostates and 3 “large” (≥90 grams) late-stage symptomatic prostates from aged patients were combined with immune cell scRNA-Seq data from 3 normal prostates from young (age 18 to 31) organ donors generated in a separate previously published study^35^. These data were analyzed together to identify common immune cell subtypes among all samples^33,34^.

Unsupervised clustering separated immune cells into 11 clusters based on immune cell gene expression profiles (Fig.1A)^33,34^. Overall, the bulk of the combined immune cells consisted of multiple closely associated T cells and NK cell clusters (Fig.1A). This is generally consistent with previous studies, which identified T lymphocytes as the predominate immune cell type in the prostate^15,24,33,34,36–38^. Since T cells and NK cells were the predominant immune cell type in the combined sample types, we first investigated alterations in these populations between normal and aged prostates. Cells from clusters identified as T and NK cells in the initial clustering analysis were combined and subclustered together, which separated the combined T and NK cells into 12 subclusters (Fig.1B, 1C, S1A, S1B). Subclusters were subsequently identified based on a comparison of gene expression profiles to those previously generated by ProjecTILs (Fig.1D, S1C)^39^. A CD8^+^ subcluster identified as a central memory (T_CM_) subtype (subcluster 0) made up the largest proportion of T cells in normal prostates (Fig.1B,1C,1D). While the overall number of immune cells was increased in aged prostates, the proportion of subcluster 0 T_CM_ cells was significantly decreased in both large and small aged prostates compared to young normal prostates (Fig.1C, S1D). A CD8^+^ subcluster (subcluster 4) with high expression of *GZMK* and low *GZMB* and an effector memory (T_EM_)-like gene profile that was proportionally significantly increased in aged versus young normal prostates was also identified (Fig.1B-E, S1B-D)^2^. These data suggest a shift in CD8^+^ T cell populations in aged prostates. Subcluster 4 was of particular interest due to its similar gene expression to a T cell subset previously associated with aging and referred to as Taa cells^2^, prompting further investigation of this subset and its potential relationship with age-related inflammation and BPH symptoms.

**Fig 1.**
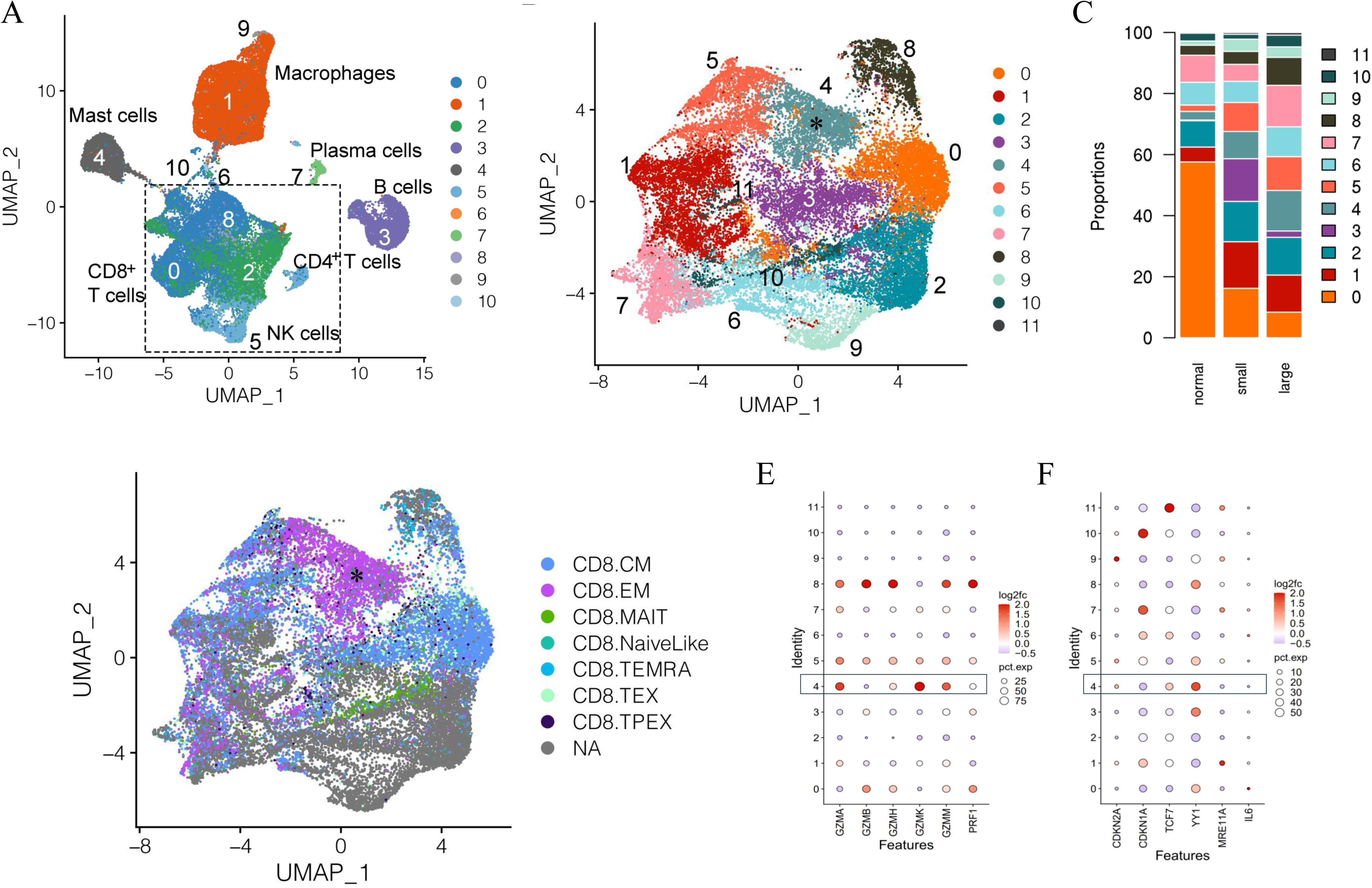
Clustering and subclustering of combined normal and aged prostate immune cells. A) Full clustering of all combined immune cells from normal young (N=3) and aged (N=13) prostates showing T/NK cells as the dominant combined immune cell subtypes. B) Subclustering of T/NK clusters identifying 12 subclusters. Subcluster 4 (asterisk) highly expresses *GZMK*. C) Proportions of T/NK cell subclusters in young normal, small, and large prostates. D) Identification of CD8^+^ T cell subsets among the subclustered T/NK cells based on ProjecTIL gene profiles; cells identified as CD8^+^ T_EM_ (asterisk) corresponding to subcluster 4, highly express *GZMK.* E) Cytotoxic protein expression of T/NK subclusters showing higher *GZMK* and lower *GZMB* expression in subcluster 4 cells from aged prostates compared to young normal prostates. F) Expression of select senescence-associated genes among subclusters in aged compared to normal prostates.

In addition to a significant increase in *GZMK* expression and decrease in *GZMB* expression, the subcluster 4 Taa cells from aged prostates also showed relatively low expression of the perforin gene *PRF1* (Fig.1E). This cytotoxic protein expression pattern has been previously associated with an early-stage CD8^+^ memory T cell differentiation state with diminished capacity for cytotoxic activity^30,40^. Expression of select cell cycle and T cell differentiation genes previously reported to be modulated in aged immune cells were differentially expressed in aged Taa cells (Fig.1F)^41–44^. Expression of *YY1*, a transcription factor linked to effector T cell differentiation, autoimmunity, and senescence was significantly increased in aged prostate Taa cells (Fig 1F)^43^. Expression of *CDKN2A* and *TCF7*, which encode the p16 cell cycle regulator and the T cell factor 1 (TCF1) protein associated with T cell development and maturation, respectively, were also mildly increased in aged Taa cells (Fig.1F)^41–43^. Overall, these results indicate alterations in the proportion and cycle-associated gene expression profiles between Taa in young and aged prostates.

### The Taa subset is clinically correlated with LUTS in BPH prostates

We hypothesized that the Taa or other T cell subsets would be correlated with BPH and LUTS in aged patients. We compared immune cell scRNA-Seq generated from aged small and large prostates^33,34^. To enhance the analysis of the Taa subset, additional immune cell scRNA-Seq data from seven large prostates were generated and integrated with the scRNA-Seq data from the thirteen aged prostates for a total of 10 small and 10 large prostate samples^33,34^.

The bulk of immune cells in the combined large and small aged prostates consisted of multiple closely associated clusters identified as T cells and NK cells (Fig.2A, Fig.S2A)^33,34^. T/NK cell subclustering analysis identified 14 subclusters (Fig.2A,S2A). Subcluster 1, identified as a CD8^+^ T_EM_-like subtype based on comparison of its gene expression profile to ProjecTILs profiles, expressed a similar high *GZMK* and low *GZMB* expression pattern as the Taa cells identified in the previous analysis (Fig.2B,S2A)^2,39^. A correlation analysis revealed that subcluster 1 Taa (R=0.7, P=0.0416) was significantly positively correlated with IPSS scores (Fig.2C)^33,39^. Additionally, a CD8^+^ subset identified as T_MAIT_ (subcluster 5) (R=0.5, P=0.0241) and a mixed T/NK cell subset (subcluster 13) (R=0.3, P=0.0282) were also positively correlated with IPSS (Fig.2B,2C). Given this correlation of Taa with IPSS as well as previous aging studies on Taa cells, we focused our subsequent studies on Taa cells.

**Fig 2.**
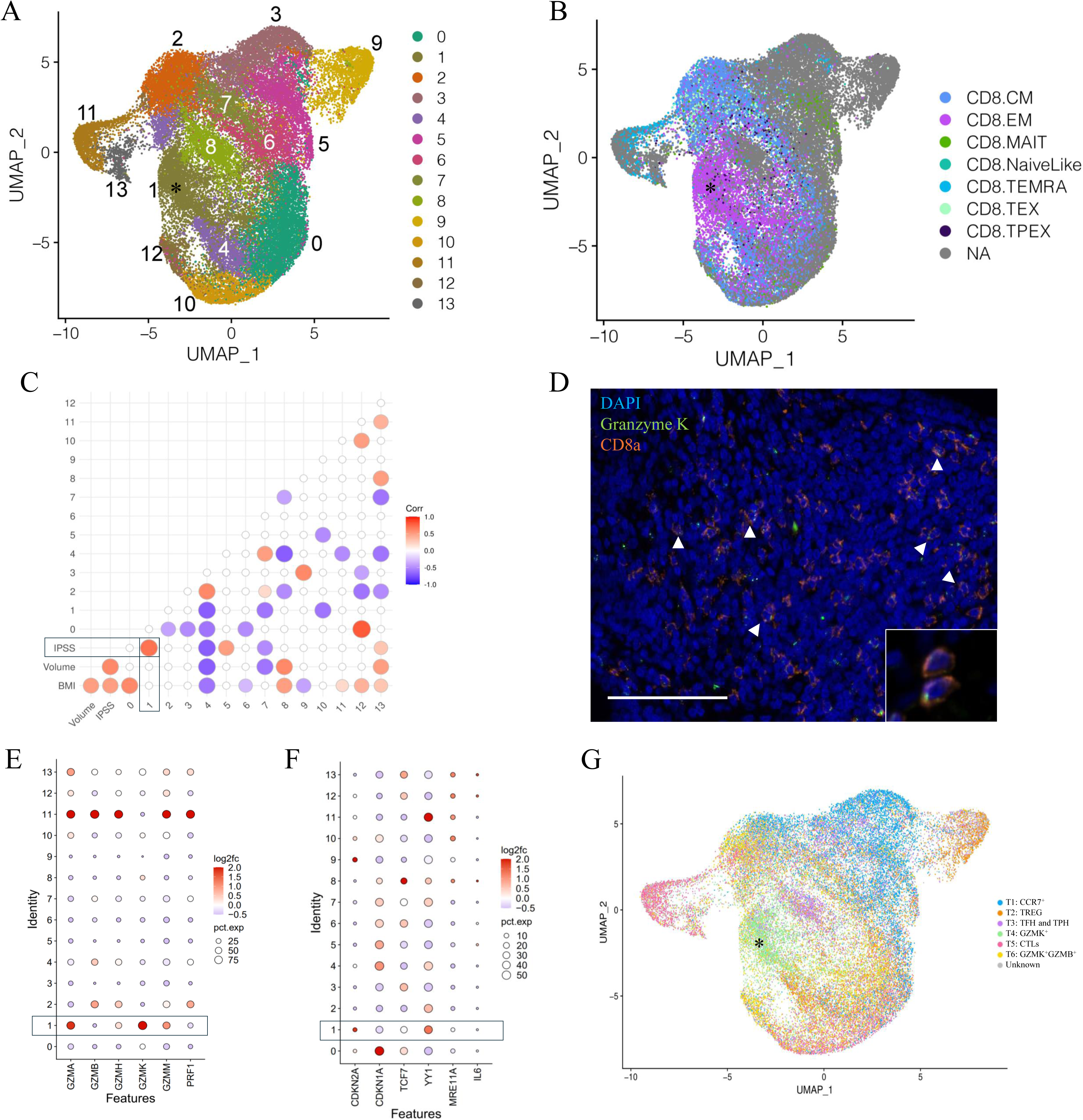
Clustering and subclustering of combined large and small aged prostate immune cells. A) Figure adapted from Lanman et al (2024)^33^. Subclustering of T/NK clusters identifying 14 T/NK cell subclusters. A *GZMK*^hi^*GZMB*^low^ subcluster designated Taa (asterisk) is identified. B) Identification of CD8^+^ T cell subsets among the subclustered T/NK cells based on ProjecTILs gene profiles; subcluster 1 Taa (asterisk) with a CD8^+^ EM-like gene expression profile, highly expresses *GZMK*. C) Correlation of IPSS, prostate volume, and BMI with T/NK cell subsets; statistically significant (P<0.05) correlations are shown. D) CD8a and GZMK-labeling T cells (arrowheads, inset) are identified by immunofluorescence within the prostate stromal compartment. Scale bar=100um. E) Granzyme expression of T/NK subclusters showing increased *GZMK* and decreased *GZMB* expression in subcluster 1 cells from large aged prostates compared to small aged prostates. F) Expression of select senescence-associated genes among subclusters in aged compared to normal prostates. G) Comparison of aged prostate and RA-derived T cells showing a common *GZMK*^+^ subcluster (asterisk). TREG: regulatory T cells, TFH: Follicular helper T cells, TPH: Peripheral helper T cells, CTL: Cytotoxic T lymphocytes.

Lymphoid cells expressing both CD8a and GZMK were observed histologically in both small and large prostate specimens, predominately within the stroma (Fig. 2D). The percentage of CD8a^+^ cells that also labeled with GZMK was increased, albeit not significantly (P=0.0614), in large prostate tissue sections (Fig.S2B). Similarly, the proportion of the Taa (subcluster 1) cells was not significantly increased in large prostates compared to small prostates in the scRNA-Seq analysis (Fig. S2C, S2D). However, there was a mild but significant increase in *GZMK* gene expression and decrease in *GZMB* gene expression as well as overall low gene expression of *PRF1* in this subcluster in large prostates compared to small (Fig.2E). Consistent with the comparison of aged and young prostate Taa cells, *YY1* and *CDKN2A* gene expression was significantly increased in Taa cells from large prostates compared to small prostates; however, *TCF7* expression was not significantly different (Fig. 2F).

Given the previous identification of *GZMK*-expressing T cells within inflamed synovial tissues of rheumatoid arthritis (RA) patients^45,46^ and the drastically increased incidence of BPH among RA patients^34,45^, we hypothesized that similar *GZMK*-expressing subsets would be present in both RA and aged prostate tissues, indicating a similar pro-inflammatory tissue microenvironment. Comparison of prostate T cell subclustering data with T cell subclustering profiles generated by Zhang et al (2019)^45^ from human RA patient synovium identified 6 similar T cell subsets including a common *GZMK*-expressing subset corresponding to the Taa subcluster (Fig.2G)^45^. In all, these results support a pro-inflammatory role for *GZMK*-expressing T cells in BPH and a potential link between age-associated T cells and symptomatic BPH^45,46^. Also, the stromal location of the CD8a and GZMK-labeling cells may suggest interactions with stromal cells contribute to BPH.

### T cell differentiation is shifted to the Taa subset in BPH prostates

Previous studies have described the association between CD8^+^ memory T cell cytotoxic granules and T cell differentiation^30^. *GZMK*^+^*GZMB*^-^ expression has been associated with early differentiation, *GZMK*^+^*GZMB*^+^ with intermediate differentiation, and *GZMK*^-^*GZMB*^+^ with late-stage differentiation^30^. We hypothesized that the differentiation trajectory would be altered in large prostate CD8^+^ T cells compared to small prostates in the direction of the GZMK^hi^GZMB^low^ Taa population. We applied a velocity analysis to the CD8^+^ T cell subclusters to analyze the differentiation trajectories of the subclusters^47^.

In both small and large prostates, overall directional flow indicated movement through CD8^+^ T_CM_ subclusters 12, 10, and 0 with most cells transitioning from T_CM_ subcluster 0 towards T_CM_ subcluster 7 (Fig3A,3B). In small prostates, the proportion of T_CM_ subcluster 7 cells is significantly increased compared to large prostates, and most of these cells progress towards T_CM_ subcluster 2 which is also significantly increased compared to large prostates (Fig.S2). T_CM_ subcluster 2 expresses both *GZMB* and *PRF1* (Fig.2D), a profile associated with cytotoxicity^30^. In contrast, overall movement of T cells from large prostates trended away from T_CM_ subcluster 2 and towards having gene expression profiles similar to the Taa subcluster 1 (Fig.3B). Further cell cycle analyses indicated that the Taa cells were cycling with high G2M scores, particularly in large prostates, suggesting proliferation within this population (Fig. S3A, S3B). Overall, these results suggest altered directional flow and differentiation of the CD8^+^ T cell subclusters between large and small specimens. This altered differentiation may contribute to the overall change in T cell subcluster proportions between small and large prostates through diversion of trajectory from a *GZMB* and *PRF1*-expressing cytotoxic population (T_CM_ subcluster 7) towards the proliferating Taa as a terminal phenotype in large prostates. Notably, subcluster 7 is also statistically significantly negatively correlated (R=-0.5, P=0.0294) with IPSS (Fig.2C), further suggesting that this shifted trajectory contributes to LUTS.

**Fig 3.**
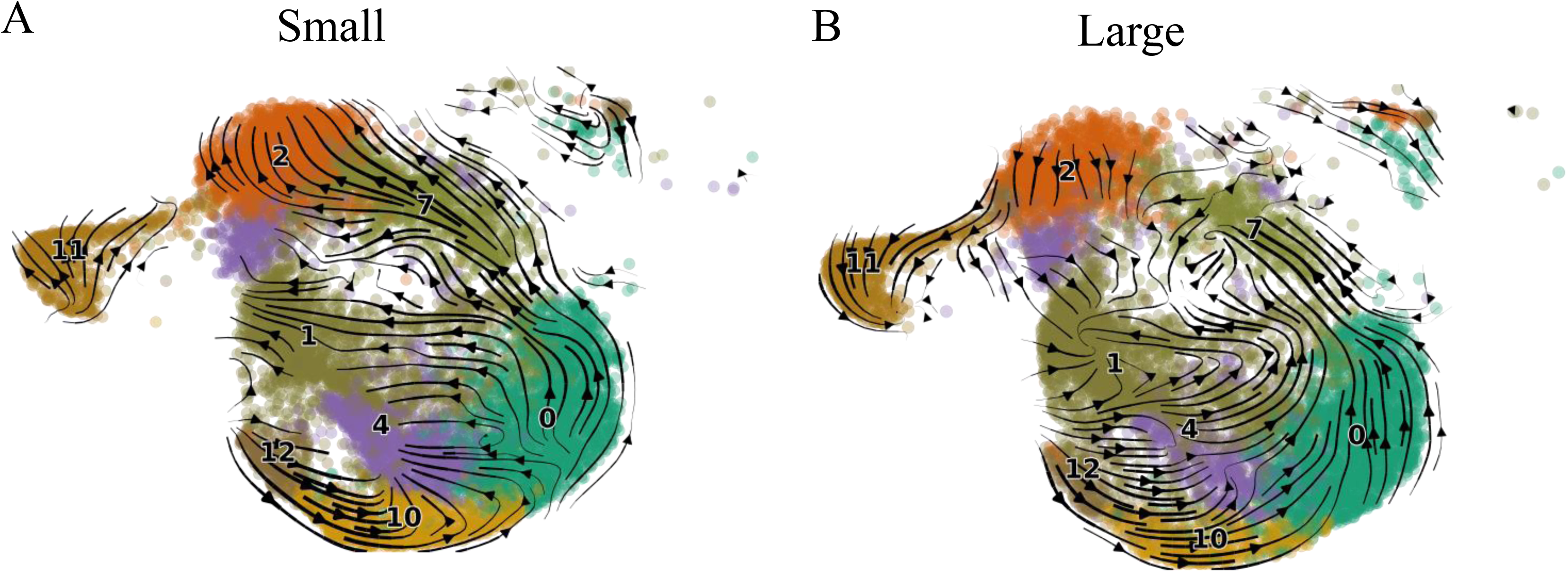
Velocity analysis of small and large aged prostate CD8 T cell subsets. A) Velocity results for small prostate CD8^+^ T cells. B) Velocity results for large prostate CD8^+^ T cells. Arrows indicate directional flow.

### SASP and senescence-associated genes are expressed by BPH stromal cells

As the Taa cells were located predominantly within the stromal compartment, and that granzyme K has been shown to induce SASP in stromal cells in other organs^2,27^, we hypothesized that these cells might impact SASP production by stromal cells and contribute to a proinflammatory microenvironment^2,27^. Using combined all-cell scRNA-Seq data generated from 5 large BPH prostates^34^, we subclustered the stromal cells and identified genes associated with SASP and senescence, and particularly expression of SASP-associated cytokine genes reportedly induced by granzyme K^2,27^. We also subclustered T cells to determine whether the Taa subcluster identified in the immune cell analyses could be identified in the all-cell analysis.

Stromal cell subclustering identified 8 subclusters segregated into two groupings of 4 subclusters, with one grouping identified as smooth muscle cell/myofibroblasts and the other as fibroblasts based on DEGs (Fig.4A). Expression of SASP-associated genes among stromal cell subclusters showed variable expression among the subclusters, including genes for GZMK-induced cytokines *CXCL8*, *CXCL1, IL6*, *CCL2*, and *CXCL2* (Fig.4B)^2,27^. Additionally, a *GZMK*^hi^ *GZMB*^low^ Taa subcluster was identified among T cells (Fig. S4A, S4B). These results indicate SASP-associated pro-inflammatory cytokine gene expression by prostate stromal cells.

**Fig 4.**
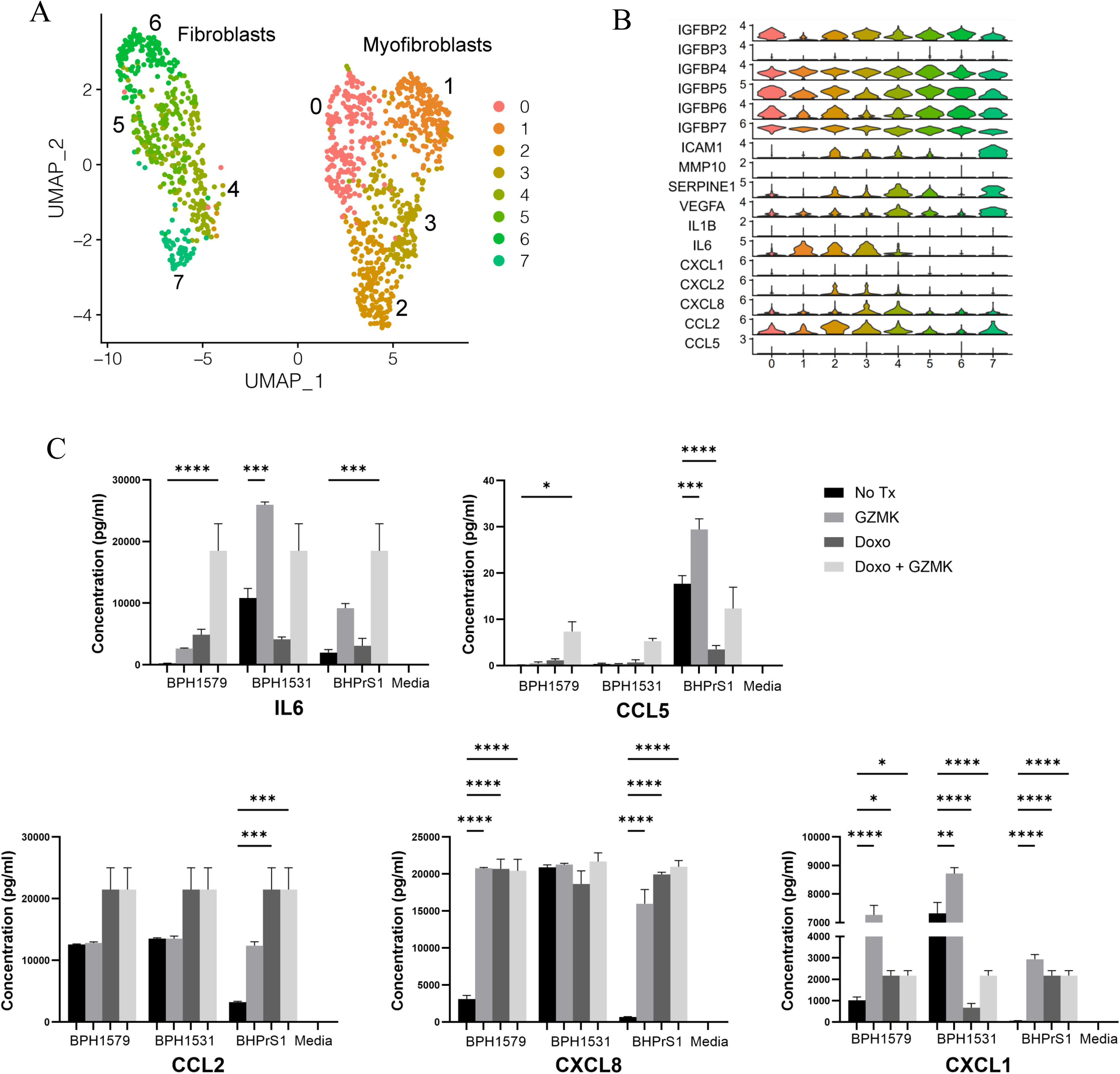
SASP-associated cytokine and chemokine expression among BPH prostate stromal cells A) Subclustering of stromal cells from 5 large aged prostates B) SASP-associated gene expression among stromal cell subclusters. C) Production of SASP-associated cytokines with at least one statistically significantly (P<0.05) modulated specimen following in vitro granzyme K treatment of cycling and Doxorubicin (Doxo)-treated senescent BPH patient-derived fibroblasts. Adjusted P values for all comparisons are listed in Supplemental Table 1.

### Granzyme K modulates SASP-associated cytokine production by cycling and senescent BPH fibroblasts

Since scRNA-seq data showed senescence-associated and SASP-associated cytokine/chemokine gene expression in prostate stromal cells, we hypothesized that granzyme K secreted by Taa may modulate senescence-associated gene expression in fibroblasts and stimulate BPH fibroblasts to produce SASP-associated cytokines/chemokines. To assess the potential impact of granzyme K on prostate stromal cells, BPH patient-derived fibroblast cell line BHPrS1 and primary fibroblasts derived from two BPH patients^48^ were treated with granzyme K in vitro. Doxorubicin (Doxo) treatment to induce senescence ^49^ was included to observe whether granzyme K stimulation can further promote SASP in senescent fibroblasts.

In both cycling and Doxo-treated senescent fibroblasts, in vitro granzyme K treatment significantly increased (P<0.05) secretion of one or more SASP-associated cytokines IL-6, CXCL8, CCL5, CCL2, and CXCL1 in at least one prostate fibroblast specimen (Fig.4C, Supplemental Table 1), and this effect varied among the fibroblast specimens. Overall, these data show that granzyme K can modulate SASP-associated factor secretion by BPH-derived fibroblasts and suggests that granzyme K may impact the prostate inflammatory microenvironment by promoting inflammatory cytokine production by both cycling and senescent prostate stromal fibroblasts. Additionally, the variability in granzyme K-induced cytokine production among the BPH patient-derived fibroblasts is compatible with the clinical and morphologic variability observed among BPH patients.

## Discussion

In this study, we identified CD8^+^*GZMK*^hi^*GZMB*^low^ Taa subset previously associated with aging present within the stromal compartment and correlated with IPSS in aged prostates^2^. Consistent with involvement in other inflammatory diseases, prostate Taa cells shared a similar gene expression profile to a *GZMK*-expressing RA T cell subset, further suggesting a potential role for granzyme K-expressing cells in promoting a dysregulated and pro-inflammatory immune microenvironment^45,46^. In vitro granzyme K treatment of prostate fibroblasts indicates granzyme K can stimulate pro-inflammatory cytokine and chemokine production by prostate stromal cells. While a specific function for Taa cells has not yet been defined^2^, these data suggest that Taa may contribute to BPH-associated inflammation and IPSS through granzyme K-mediated stimulation of stromal cell cytokine and chemokine production leading to recruitment and activation of additional immune cells in the prostate microenvironment.

Of the five granzymes (A, B, H, K, M) identified in humans, granzyme K is the least well studied ^50^. While the functions of granzyme K are currently not well defined, previous studies have identified intracellular and extracellular roles for granzyme K which include both cytotoxic and non-cytotoxic activities^50,51^. However, unlike the most common cytotoxic granzyme GZMB, which in combination with perforin (PRF1) is associated with induction of apoptotic cell death, GZMK lacks lytic activity and its expression has not been positively correlated with cytotoxic activity^30,50^. Also, unlike other granzymes, a cellular receptor for GZMK has not been identified. In addition to low *GZMB* and high *GZMK* gene expression, Taa cells showed low *PRF1* expression, further suggesting reduced cytotoxic capacity in these cells (Fig.1F,2C). T cell cytotoxic capacity and ability to eliminate infected or neoplastic cells diminishes with age, suggesting an association between an increased proportion of *GZMK*^hi^ T cells and diminished cytotoxic activity with aging^30^. Additionally, expression of *YY1*, a gene that has been associated with both cellular senescence and T cell-mediated diseases, and *CDKN2A*, a gene involved in senescence-associated cell cycle arrest, were increased in large prostate Taa cells, further suggesting alterations in this T cell population between small and large prostates^41–43^.

As T cell differentiation involves the maturation of lytic granules, we hypothesized that the changes in granzyme gene expression may indicate altered T cell differentiation between large and small aged prostates^30,31^. RNA velocity analysis was used to infer the differentiation fates of CD8^+^ T cell subsets^47,52,53^. A high proportion of unspliced transcripts for a particular gene indicates upregulation of that gene, and RNA velocity analysis describes changes in the abundance of mRNA transcripts^47^. By examining the velocities of multiple genes, the differentiation pathways of individual cells may be inferred^47^. Genes were ranked on velocity to find genes in a subcluster that show dynamics that are transcriptionally differently regulated compared to all other subclusters, for example, induction in that cluster and homeostasis in the remaining population. In the current study, directional flow to and from the Taa population is altered between large and small aged prostates. ^30^. Also, evidence of cycling in the Taa subcluster suggests these cells are proliferating, which may also contribute to the accumulation of these cells in large prostates.

Overall, findings in the current study suggest a potential role for the age-associated Taa in BPH. It is hypothesized that the interplay of Taa cells and stromal cells may contribute to the non-resolving and progressive inflammation of BPH though granzyme K-mediated SASP cytokines and chemokines. A similar effect has been described in concurrent studies involving myeloid cells^33^. However, while the secretion of multiple SASP cytokines and chemokines were modulated by prostate fibroblast granzyme K treatment in vitro, further studies are needed to confirm a direct fibroblast stimulatory capacity for Taa cells and if Taa cells have a causal role in the development and progression of BPH-associated inflammation and LUTS. Additionally, further study is needed to determine the underlying mechanisms for altered T cell differentiation and gene expression and to define the clinical consequences of these alterations. For example, in addition to SASP genes, stromal cell subclusters expressed *CDKN1A*, *YY1,* and *IL6*, genes that have been previously linked to cellular senescence (Fig.S4C)^54^. Additionally, YY1, which has been shown to repress p16 expression and curtail senescence in cancer cells, has also been associated with collagen production and implicated in fibrotic diseases such as idiopathic pulmonary fibrosis (IPF) and liver fibrosis through promoting myofibroblasts differentiation and increased collagen production by fibroblasts and myofibroblasts^55–58^. As other studies have suggested a role for prostatic fibrosis in contributing to BPH-associated LUTS, these results may indicate a role for YY1 signaling in BPH^59^. More recently, a role for granzyme K-mediated complement activation in RA and other chronic inflammatory conditions has been described^60^. Complement has been shown to be highly expressed by synovial fibroblasts and promote fibroblast-mediated inflammatory tissue priming in RA^61^. Given the similarities between BPH and RA inflammation, granzyme K-mediated complement activation and fibroblast stimulation may play a role in driving BPH inflammation and LUTS.

While the current study focused on CD8^+^ T cells and specifically Taa cells, other T cell subsets were positively correlated with IPSS and/or prostate volume (Fig.2C). This includes a CD8^+^ T_MAIT_ subset, and CD8^+^ T_MAIT_ cells have previously been implicated in immune-mediated inflammatory diseases^62,63^. Future studies of the role of aging and T cells in BPH include investigating this and other specific T cell subsets, as well as epigenetic alterations or changes in T cell receptor (TCR) repertoires with age, and exploring interactions between stromal and epithelial cells with immune _cells11,64,65._

### Limitations of the study

Several challenges were encountered in this study. As surgical intervention is not indicated for patients without bothersome urinary symptoms or without evidence of PCa, therefore small, aged prostates were collected from PCa patients with small PZ-confined tumors^33,34^. The field effects of PCa tumors are considered limited to within about 3mm of the tumor periphery^66,67^. For these reasons, PCa-free TZ tissue obtained from these specimens is an accepted method for collection of these samples^66,67^. However, the potential effects of PZ tumor cells on TZ immune cells cannot be completely ruled out. Another challenge in this study was related to the previously published normal non-BPH prostate scRNA-Seq study by Henry et al. (2018), which all prostate cells rather than isolated CD45^+^ leukocytes were analyzed^35^. The inclusion of more numerous prostate cell populations along with the relatively small number of immune cells normally present in non-BPH prostates meant there were relatively few normal prostate immune cells for analysis compared to our BPH specimens. Consequently, this may have affected some analyses and prevented the inclusion of normal immune cells in some analyses.

## Acknowledgments

The authors thank Dr. Philip San Miguel of the Purdue Genomics Core. We acknowledge the assistance of Dr. Abigail Cox, MacKenzie McIntosh, Megan Cohen, and Victor Bernal-Crespo of the Purdue University Histology Research Laboratory, a core facility of the NIH-funded Indiana Clinical and Translational Science Institute with the preparation of histological and immunofluorescence sections. We gratefully acknowledge the support of the Purdue University Genomics Core Facility, the Collaborative Core for Cancer Bioinformatics (C3B), which is funded by the Walther Cancer Foundation and the Purdue University Institute for Cancer Research (NIH grant P30 CA023168), and Purdue Research Computing for their continued support. This work was funded by 5P20DK116185 from NIDDK.

## Author contributions

APG and BTH collected surgical prostate specimens. REV and GC performed the aged prostate tissue processing for scRNA-Seq. NAL, HK, JSPP, and AK performed the scRNA-Seq analyses. MMB, REV, and GC analyzed the resulting scRNA-Seq gene expression data. MMB designed and conducted the in vitro experiments and performed the histologic evaluation. MMB and NAL wrote the manuscript text. GH and DWS provided the previously generated normal prostate scRNA-Seq data. This work was funded through 1P20DK116185 from NIDDK.

## Methods

### Human prostate samples

Prostatic tissues were obtained as described in Vickman et al (2022), Lanman et al (2024), and Henry et al (2018) ^33–35^. Clinical data for BPH patients is summarized in Lanman et al (2024)^33^. All human tissue procurement was done in accordance with protocols approved by the Institutional Review Boards of each institution.

### BPH tissue processing for scRNA-Seq

Normal prostate tissues were processed for scRNA-Seq as described in Henry et al (2018)^35^. For aged prostate tissues, transitional zone (TZ) tissue was excised from each collected large and small prostate and processed as described in Vickman et al (2022) and Lanman et al (2024) and in Supplemental Methods SM1^33,34^.

### Sample sequencing and data analysis

Sequencing of normal prostate cells was performed as described in Henry et al (2018) and is accessible through GEO SuperSeries GSE120716 ^35^.

Sequencing of aged prostate cells was performed as described in Vickman et al (2022), Lanman et al (2024) and Supplemental Methods SM1^33,34^. The aged prostate data is available through GEO under accession number GSE269205. For comparison of aged prostate to normal young prostate immune cells, published scRNA-seq data generated from immune cells isolated from 13 aged prostate specimens was combined with previously published scRNA-seq data generated from 3 normal young prostates^35^. For comparison of large and small aged prostate immune cells, published CD45^+^ leukocyte scRNA-seq data from the 10 large and 10 small aged prostates were used. Clustering and subclustering of cells was performed as described in Vickman et al (2022) and Lanman et al (2024) and Supplemental Methods SM1^33,34^.

### Clinical correlation

Cellular proportions for each cluster and subcluster were computed for each sample. A permutation test was used to calculate a p-value for each cluster, utilizing bootstrapping to calculate the confidence interval for the magnitude of difference between large and small prostates from aged men and young normal prostates. Spearman correlations were computed and statistically significant correlations identified (adjusted p-value < 0.05) between cellular proportions and patient body mass index (BMI), IPSS, and prostate volume using the corrplot R package version 0.92.

### Velocity analysis

The package Velocyto^68^ v. 0.17.17 was used to count spliced and unspliced abundances from the CellRanger output BAM files and write these abundances to loom files. The metadata, which includes sample numbers, genes post-filtering, cells post-filtering, embedding coordinates of cells, and clusters were output from Seurat^69^ v3 and used in the RNA velocity analysis. Next, scVelo^47^ v. 2.4.0 was used to estimate RNA velocity using a likelihood-dynamical model. This dynamical model allows the estimation of RNA velocity even if there is not steady state observed for a given gene, provided that enough spliced and unspliced counts are observed.

### Prostate immunofluorescence

Full thickness cross sections of prostate were fixed in 10% neutral buffered formalin (NBF) for histology. Formalin fixed prostate tissues were embedded in paraffin and sectioned. Sections were deparaffinized and blocked with 2.5% Normal Goat serum for 20 minutes. Sections were incubated with primary antibodies for CD8a (clone C8/144B, Abcam) at 1:100 and Granzyme K (polyclonal, Novus Biologicals) at 1:50 for 60 minutes the rinsed twice with staining buffer. The sections were incubated with GoRb488 at 1:250 and GoM555 at 1:500 for 30 minutes then rinsed once with buffer. Sections were counterstained with DAPI for 15 minutes and rinsed with water before applying Prolong Gold mounting media and coverslipping. Stained slides were digitized using a Leica Versa8 whole-slide scanner (Leica, Wetzlar, Germany) and analyzed using the Visiopharm digital slide analysis platform (Visiopharm, Hørsholm, Denmark).

### BPH prostate fibroblast and T cell scRNA-Seq analysis

Previously published scRNA-seq data generated from all viable cells following tissue digests of 5 large prostate TZ obtained from patients undergoing simple prostatectomy surgery for symptomatic BPH was used to analyze BPH T cells and fibroblasts^34^. Both raw and processed scRNA-seq data are available on GEO under accession number GSE183676.

### Rheumatoid arthritis T cell analysis

To compare the BPH T cell subset to rheumatoid arthritis-derived T cell subsets, we trained a classifier based on T cell subset marker genes from the supplemental data from Zhang et al, (2019)^45^. The top 20 genes identified for the T cell subsets CCR7+(SC-T1), Treg (SC-T2), Tph and Tfh (SC-T3), GZMK^+^(SC-T4), CTLs (SC-T5), and GZMK^+^GZMB^+^ (SC-T6) T cells were used as input into Garnett^45,70^. The Garnett classifier was trained using default parameters and then classified our cells with the resulting model using default parameters.

### Human BPH fibroblast culture and cytokine analysis

Human prostate fibroblasts were obtained from the freshly isolated transition zones of 2 BPH patient specimens following simple prostatectomy surgery and isolated as previously described^34^. Isolated prostate fibroblasts along with the human prostate fibroblast cell line BHPrS1 were plated at a density of 10,000 cells per well in a 96 well tissue culture plate in complete RPMI + 10% FBS^48^. Cells were grown to around 70% confluency then serum starved in RPMI + 0.1% FBS for 24 hours, then treated with 200nM recombinant granzyme K (MyBioSource, San Diego, CA) in RPMI + 1% FBS for 24 hours^27^. For senescence studies, fibroblasts were treated with 250nM doxorubicin (Sigma-Aldrich) for 24 hours, then cultured for an additional 6 days in RMPI + 10% FBS^49^ prior to granzyme K treatment as described above. Supernatants were collected and analyzed for CXCL1, CCL2, CCL5, CXCL8, IL6, TNF, CCL20, IL2, and IL12p70 using the LEGENDPlex system (Biolegend) per manufacturer protocols. Comparisons with significant P values (P<0.05) are listed in Supplemental Table 1.

### Statistical analyses

Statistical analyses for scRNA-Seq analyses are described above. For IF quantification, a two-tailed T test was performed using GraphPad Prism (version 9; GraphPad Software, San Diego, CA). For ELISA experiments, a one-way ANOVA and Šídák’s multiple comparisons test were performed using GraphPad Prism. P values less than 0.05 were considered significant.

## Supplemental Figure Legends

Fig.S1 Comparison of young normal and aged prostate immune cells. A) Distribution of T/NK cells from the combined 3 young normal and 13 aged prostates among subclusters. B) Dotplot of the top differentially expressed genes from each T/NK cell subcluster. C) Identification of all T/NK cell subclusters based on ProjecTILs gene expression profiles. D) Permutation analysis showing statistical differences in subcluster proportions between young and aged sample types. E) Gene expression of *CD8a*, *GZMK*, and *GZMB* among the T/NK subclusters.

Fig.S2 Comparison of large and small aged prostate immune cells. A) UMAP of T cell subcluster identification based on ProjecTILs gene list comparison. B) Percentage of anti-GZMK-labeling cells among anti-CD8a-labeling cells immunofluorescence sections of large (n=5) and small (n=5) prostate specimens. D) Proportion of each T/NK cell subcluster in large and small aged prostates. Subcluster 1 Taa cells (asterisk) are identified as having a CD8^+^ EM gene expression profile. E) Permutation test comparing the proportion of each T/NK subcluster between large and small prostates.

Fig S3. A) Cell proliferation scores of small prostate CD8+ T cells. B) Cell proliferation scores of large prostate CD8+ T cells

Fig.S4 Analysis of T/NK cells from all-cell large aged prostate specimens. A) Subclustering of T/NK cells. B) *GZMK* and *GZMB* gene expression among T/NK cell subclusters. C) Select senescence-associated gene expression among stromal cell subclusters.

## Supplemental Methods SM1

### BPH tissue processing for scRNA-Seq

Normal prostate tissues were processed for scRNA-Seq as described in Henry et al (2018)^35^. For aged prostate tissues, transitional zone (TZ) tissue was excised from each collected large and small prostate and processed as described in Vickman et al (2022) and Lanman et al (2024)^33,34^. In summary, tissues were digested and processed to a single-cell suspension. Samples for immune cell-only analysis were incubated with 5µl Human TruStain Fx Blocking reagent (Biolegend, San Diego, CA) and 0.5µl Zombie Viability Dye (Biolegend) in 100µl PBS per sample for 10-15 minutes at RT in the dark, filtered and spun, then incubated with an antibody cocktail of CD45-PE (clone HI30, Biolegend) pan-leukocyte marker, EpCAM-APC (clone 9C4, Biolegend) epithelial cell marker, and CD200-PE/Cy7 (clone OX-104, Biolegend) endothelial cell marker or single antibodies for compensation controls (if necessary) for 30 minutes at 4°C. Samples were washed with PBS spun down and then resuspended in complete RPMI for live cell sorting on the BD FACS ARIA II (BD Biosciences, Franklin Lakes, NJ) to isolate CD45^+^EpCAM^-^CD200^-^ immune cells. Cellular Indexing of Transcriptomes and Epitopes by Sequencing (CITE-seq) was performed on cells from 9 large and 3 small prostate tissues, allowing quantification of protein expression^34,71^. Tissues were stained with TotalSeq-B Antibodies (Biolegend) CD3 [clone UCHT1], CD4 [clone RPA-T4], CD8 [clone RPA-T8], CD11b [clone ICRF44], and CD19 [clone HIB19] per the manufacturer instructions, prior to loading into the Chromium System (10x Genomics, Pleasanton, CA). Sorted and stained immune cells from each BPH sample were counted, prepped, and loaded into the 10X chip for a 5000 target cell recovery per 10X Genomics protocols. cDNA synthesis and clean-up steps were performed per manufacturer protocols. cDNA content and quality were assessed via Agilent Bioanalyzer (Agilent Technologies, Santa Clara, CA). Sample library preparation was performed per 10X Genomics protocols prior to sequencing.

### Sample sequencing and data analysis

Sequencing of normal prostate cells was performed as described in Henry et al (2018)^35^. Sequencing of aged prostate cells was performed as described in Vickman et al (2022) and Lanman et al (2024)^33,34^. In summary, BPH samples were sequenced by the Purdue Genomics Core using a NovaSeq S4 flow cell on a NovaSeq 6000 platform (Illumina, San Diego, CA). Paired-end, 2×150 base-pair reads were sequenced to a depth of 50,000 reads per cell. TotalSeq-B antibody libraries for quantification of cell surface proteins were sequenced at a depth of 5,000 reads per cell. The estimated number of cells along with mean reads per cell and mean number of genes per cell for each sample are listed in Vickman et al (2022)^34^. Sequencing reads from the Chromium system were de-multiplexed, adapters removed, and reads were trimmed using the CellRanger pipeline v3.0.0 (10X Genomics). R version 3.5.1 and Bioconductor version 3.8 were used for all statistical and bioinformatic analyses. The Seurat toolkit for single-cell analyses, version 3.1.3 was used for data scaling and normalization, cell clustering based on gene expression, and identification of marker genes^72^.

Data were normalized using scTransform v.0.3.1 and cell cycle scoring was performed using cell cycle-related genes^73^. Scaling was performed, regressing out cell cycle scores, mitochondrial reads, and UMI counts using generalized linear models to remove heterogeneity due to these variables.

After scaling, dimensionality reduction was performed and the first 30 principal components were selected, followed by unsupervised clustering of cells. Clusters were identified using the Louvain method for community detection, as implemented in Seurat. A resolution of 0.2 was selected, which was determined via the clustree R package v 0.4.3, selecting a resolution that provides stable clusters^74^. P-values were corrected for multiple testing using the Benjamini-Hochberg method^75^. Biomarkers were considered statistically significant at a 1% false discovery rate (FDR) using the Wilcoxon rank sum test. Both raw and processed scRNA-seq data are available on GEO under accession number GSE164695.

For comparison of aged prostate to normal young prostate immune cells, scRNA-seq data generated from immune cells isolated from 13 aged prostate specimens was combined with previously published scRNA-seq data generated from 3 normal young prostates^35^. For comparison of large and small aged prostate immune cells, scRNA-seq data from the 10 large and 10 small aged prostates were used. Differentially expressed genes between sample groups were identified using the edgeR Bioconductor package, v 3.31 with an FDR cutoff of 5%.

To identify leukocyte clusters, a combination of CITE-seq and marker gene expression as well as comparisons with the Human Cell Atlas using the Bioconductor package SingleR were utilized^34,76^. T and NK cells expressing CD3 (CD3E, CD3D, or CD3G) were subsequently separated, variable genes identified, normalized, and then subjected to clustering and community detection with Seurat.

