## Supplemental Figures and Table for "Immune cell single-cell RNA sequencing analyses link an age-associated T cell subset to symptomatic benign prostatic hyperplasia"

### Supplemental Figure S1

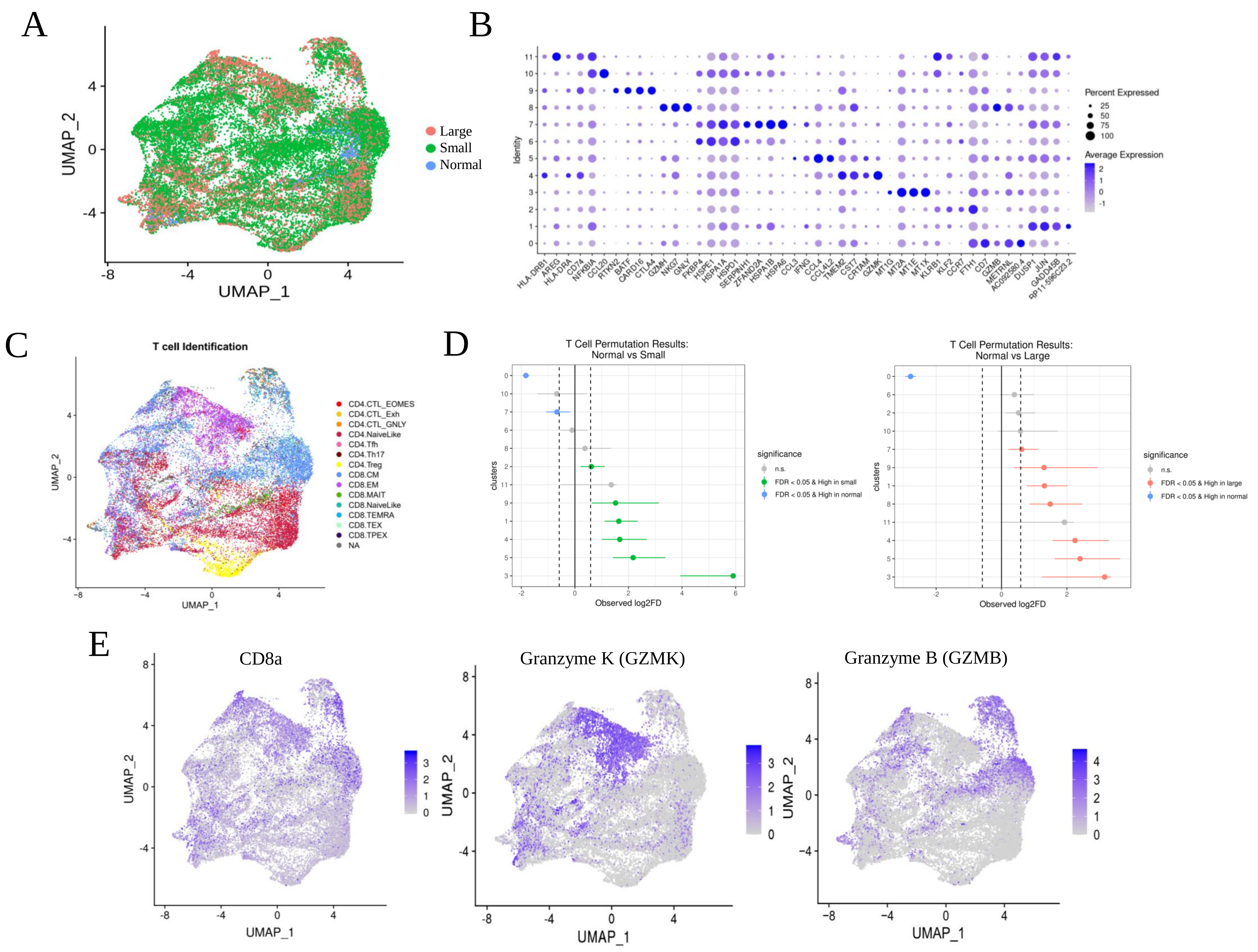

### Supplemental Figure S2

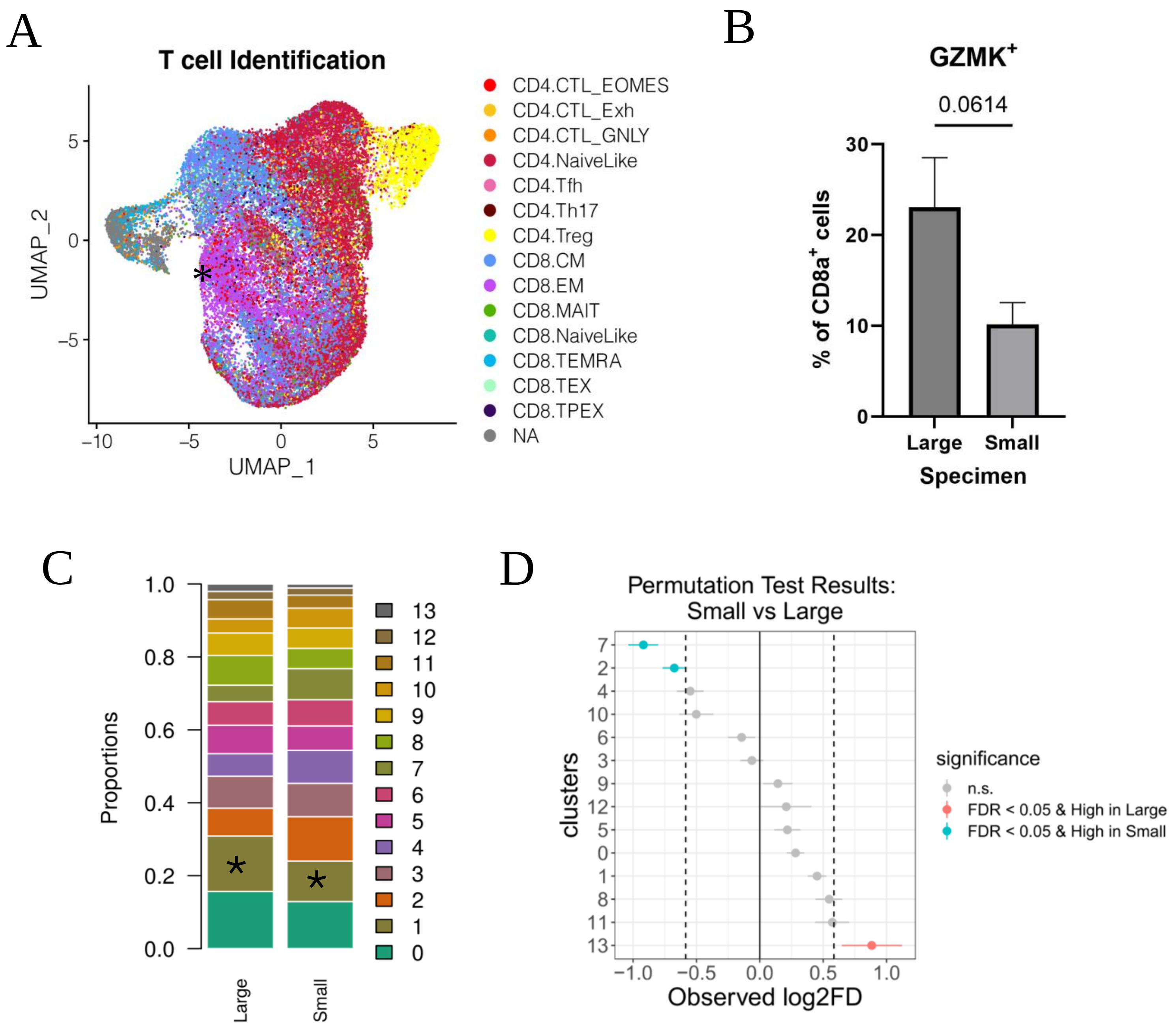

Supplemental Figure S3

A Small

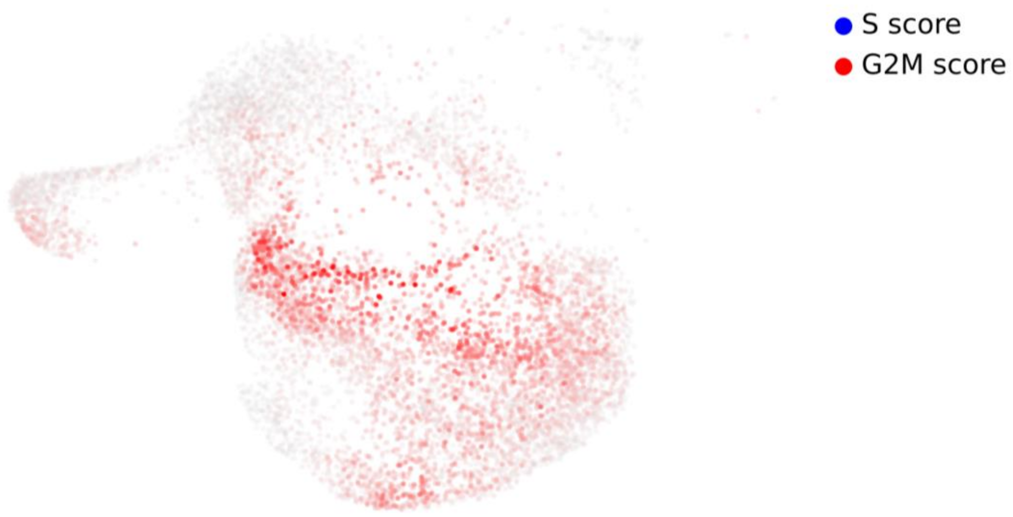


B Large

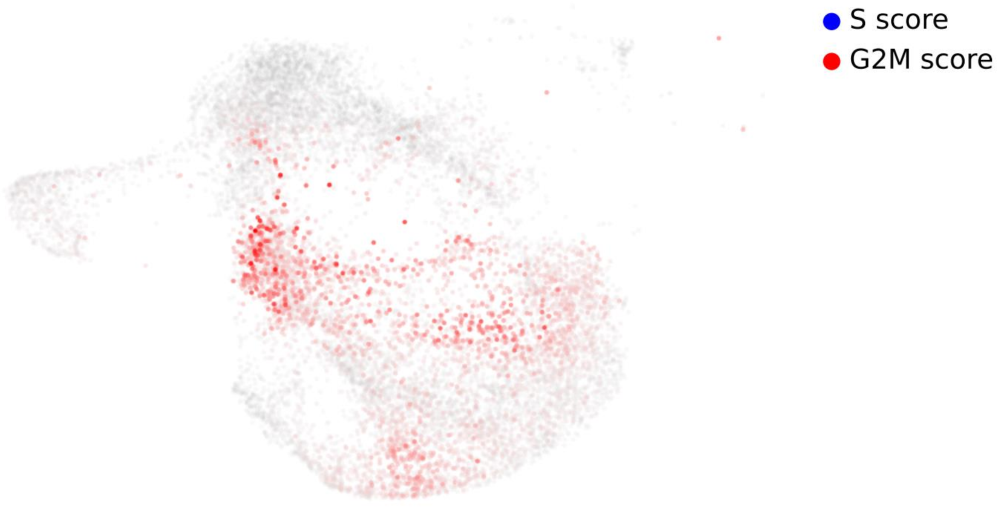

Supplemental Figure S4

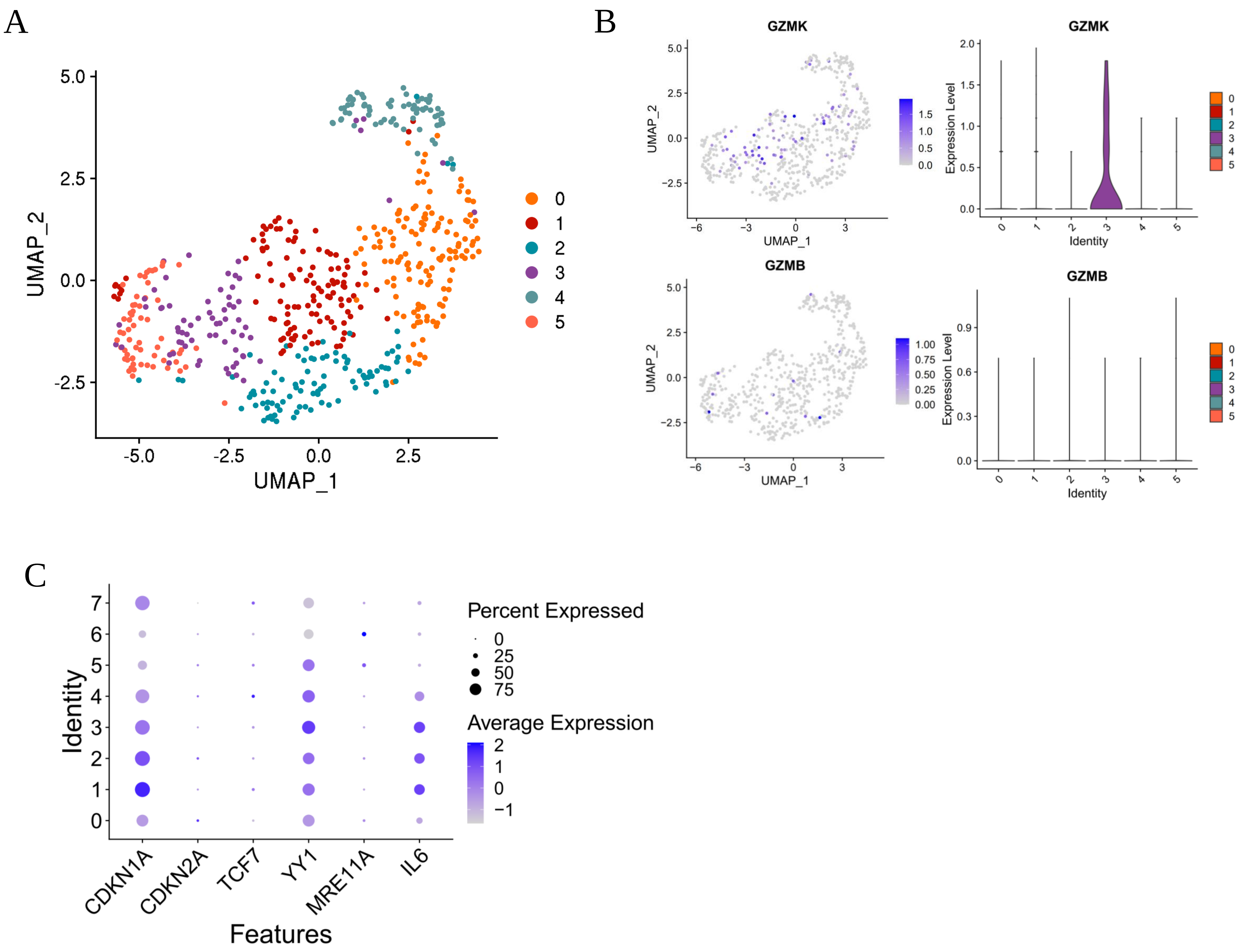
